# Protective mutation A673T as a potential gene therapy for most forms of APP Familial Alzheimer’s Disease

**DOI:** 10.1101/2020.07.22.215616

**Authors:** Antoine Guyon, Joël Rousseau, Gabriel Lamothe, Jacques P. Tremblay

## Abstract

The accumulation of plaque in the brain leads to the onset and development of Alzheimer’s disease. The Amyloid precursor protein (APP) is usually cut by α-secretase, however an abnormal cleavage profile by β-secretase (BACE1) leads to the accumulation of Aβ peptides, which forms these plaques. Numerous APP gene mutations favor plaque accumulation, causing Familial Alzheimer Disease (FAD). However, a variant of the APP gene (A673T) in Icelanders reduces BACE1 cleavage by 40 %. A library of plasmids containing APP genes with 29 FAD mutations with or without the additional A673T mutation was generated and transfected in neuroblastomas to assess the effect of this mutation on Aβ peptide production. In most cases the production of Aβ peptides was decreased by the co-dominant A673T mutation. The reduction of Aβ peptide concentrations for the London mutation (V717I) even reached the same level as A673T carriers. These results suggest that the insertion of A673T in the APP gene of genetically susceptible FAD patients may prevent the onset of, slow down, or stop the progression of the disease.

## Introduction

The ageing population in the Western world is of grave importance as both a socioeconomic issue and a strain on the medical system (1). More than 5% of the population above 60 years old is affected by dementia; of these, two thirds are the result of Alzheimer’s disease (AD) (2–4). After the age of 65, the prevalence of AD doubles every five years. As a result, individuals aged 90 years or older have a prevalence of AD of greater than 25% (4). As baby boomers, one of the most populous generational groups in American history, enter retirement, AD is becoming an increasingly heavy burden on the medical system (1, 5). As such, an increased focus on the diagnostic and treatment of this disease has become critical.

AD diagnosis is confirmed by two major histopathologic hallmarks: neurofibrillary tangles and senile plaques. The former are somatic inclusions of the microtubule-associated tau protein while the latter are comprised of extracellular deposits of amyloid β (Aβ) peptides. These plaques, comprised mostly of β-amyloid peptides, are a central pathological feature of AD (6, 7). The β-amyloid peptides are the result of sequential proteolytic processing of the amyloid-β precursor protein (APP) by β- and γ-secretases (8). APP is a membrane protein expressed in many tissues but mostly in neuron synapses.

β-secretase, also known as aspartyl protease β-site APP cleaving enzyme 1 (BACE1), preferentially cleaves APP at the β-site in exon 16 (between Met671 and Asp672) (9, 10). Subsequent cleavage by γ-secretase in exon 17 releases the β-amyloid peptides (40-42 amino acids long). In AD-free individuals, APP is preferentially processed by α-secretase prior to the cleavage by γ-secretase. The α-secretase enzyme targets the α-site located within the β-amyloid peptide sequence (Fig 1) and prevents the formation of the Aβ peptides thus reducing the formation of insoluble oligomers and protofibrils resulting from the aggregation of these peptides. These oligomers and protofibrils accumulate to form neurotoxic senile and neuritic plaques.

**Fig 1:**
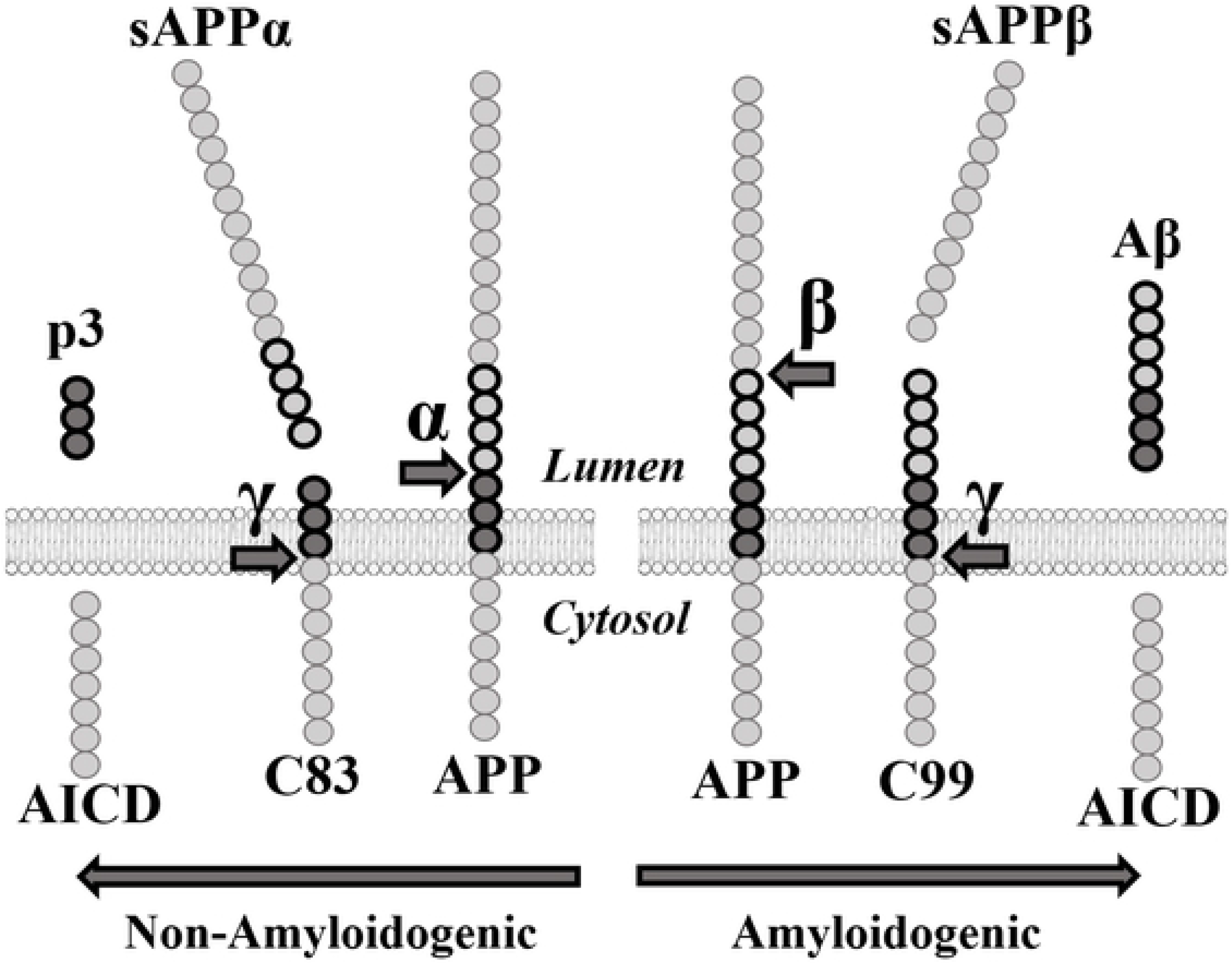
APP proteolytic pathways. **α**: α-secretase. **sAPPα:** soluble APPα fragment. **β**: β-secretase. **sAPPβ:** soluble APPβ fragment. **γ:** γ-secretase. **AICD**: APP Intracellular domain.

Initially, many big pharma companies attempted to inhibit BACE1 to decrease Aβ peptides concentration. However, numerous clinical studies targeting BACE1 failed due to notable side effects (11, 12). Indeed, BACE1 was found to be significantly implicated in several other pathways necessary for synaptic transmission (13–15). The soluble APPβ fragment (sAPPβ) generated by the cleavage of APP by BACE1 is also involved in axonal generation and neuronal death mediation (16) (Fig 1). This implies that an effective treatment for AD must decrease Aβ peptide concentrations without eliminating either sAPPβ or BACE1. Since eliminating the enzyme responsible for the excessive cleavage is unrealistic as a treatment, targeting the APP gene itself must become the focus of many future gene-based therapies.

Many APP mutations cause early-onset Familial Alzheimer’s disease (FAD). However, not all mutations are made equal; rather than causing FAD, some APP mutations decrease the incidence of this disease. Indeed, in 2012, Jonsson et al. published a study in which they searched for APP coding variants in a sample of 1,795 Icelanders using whole-genome sequencing (17). Their goal was to find low-frequency variants of the APP gene, which significantly reduced the risk of AD. Ultimately, they found a point mutation in the APP gene wherein the alanine at position 673 was substituted for a threonine (A673T). This mutation protects against AD and is adjacent to the β-site in exon 16 of the APP gene. The amino acid in question is located at position 2 in the ensuing β-amyloid peptide. Due to the proximity of the A673T mutation to the β-site, the authors proposed that A673T specifically impairs the cleavage of the APP protein by β-secretase.

In fact, the A673T mutation was shown to reduce the formation of β-amyloid peptides in wildtype APP by about 40% *in vitro* (17). The strong protective effect of the A673T mutation against AD serves as a proof of principle that reducing the β-cleavage of APP may protect against the disease. Moreover, the A673T mutation may also help to prolong the lifespan of its carriers. Indeed, individuals with this mutation were reported to have 1.47 times greater chances of reaching the age of 85 when compared to non-carriers. Jonsson et al. concluded that the A673T mutation confers a strong protection against AD (17). Later, Kero et al. found the A673T variant in an individual who passed away at the age of 104.8 years with little β-amyloid pathology (18). This report supports the hypothesis that A673T protects the brain against β-amyloid accumulation and AD.

Although the A673T mutation is protective for people with an otherwise wild type APP gene, it is not known whether the A673T mutation will reduce the formation of β-amyloid peptides when the APP gene contains a deleterious FAD mutation. We thus aimed to study the interaction between the A673T mutation and various FAD mutations to determine to what extent it can provide a protective effect in patients. If the A673T is protective for FAD, this modification could eventually be introduced in an FAD patient genome with the CRISPR/Cas9 derived gene editing technology.

We report here that the presence of the Icelandic mutation (A673T), in an APP gene containing most of the FAD mutations, reduces the production of the Aβ40 and Aβ42 peptides. The introduction of the A673T mutation could thus be an effective treatment for heritable FAD and perhaps even for sporadic AD.

## Results

### Amyloid-β peptides quantification

We first measured the concentration of Aβ peptides in the culture medium of neuroblastomas that were transfected with a wild-type APP plasmid or not (Fig 2). The Aβ42 concentration of cells not transfected with the plasmid was so low that it was below the detection range of the MSD kit. There was, however, a clear increase of Aβ40 and Aβ42 peptide concentrations for the cells transfected with the wild-type APP plasmid.

**Fig 2:**
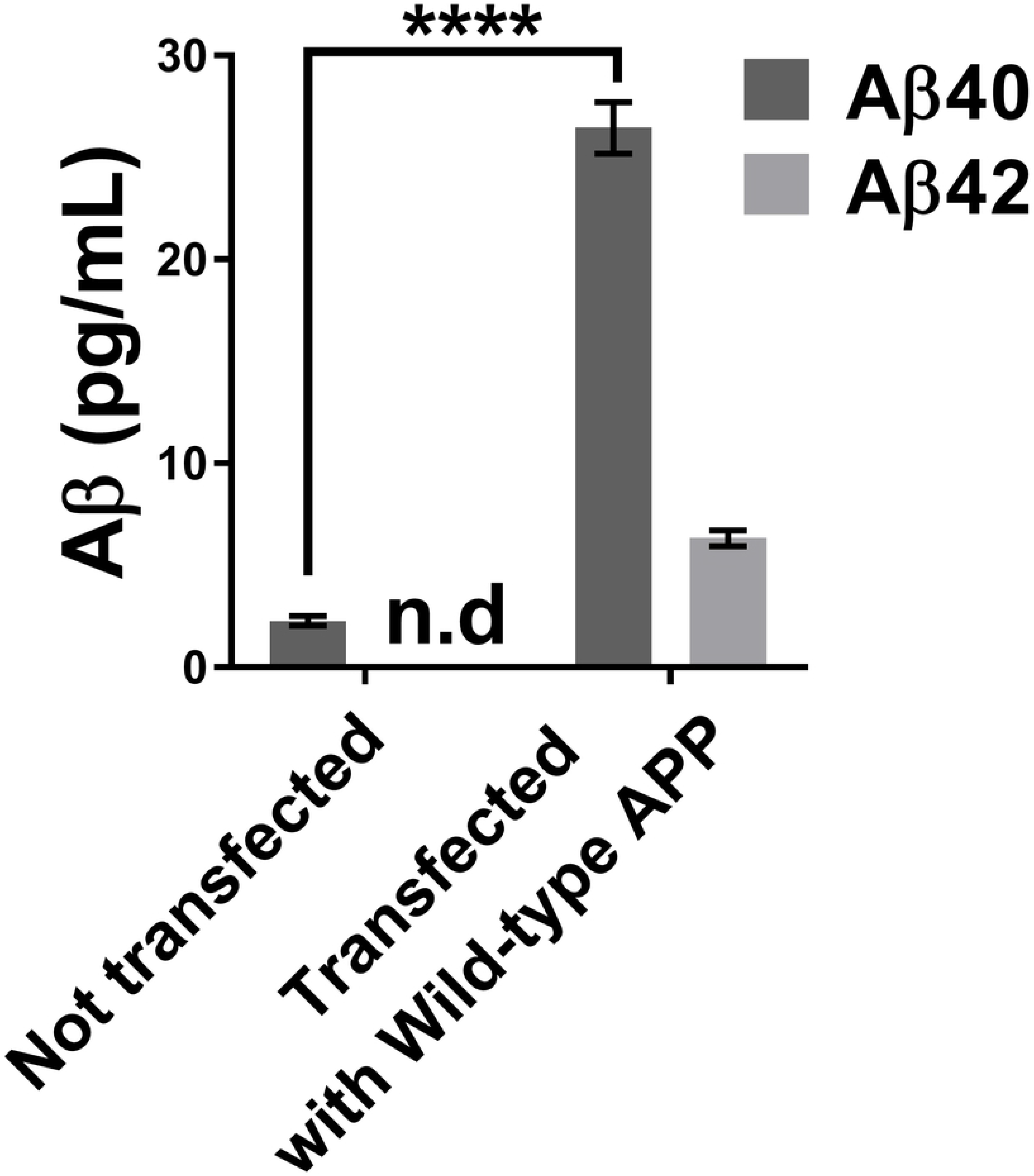
Concentrations of Aβ40 and Aβ42 in the neuroblastoma supernatant using the MSD Elisa test. The neuroblastoma transfection with a wild APP plasmid increased the concentrations of Aβ40 and Aβ42 peptides in the cell culture medium. Statistic test: Two-way ANOVA Sidak’s multiple comparisons test. P value style **** p<0.0001. n.d: not detectable

Two plasmid libraries were then tested. One library was comprised of the APP plasmid with a unique FAD mutation in each plasmid. The other was composed of the same APP/FAD plasmids but with the additional A673T mutation. We first analyzed the effect of the different FAD mutations on the Aβ peptide concentrations in the cell culture medium (Fig 3). Nearly all of the FAD mutations increased the Aβ40 and Aβ42 concentrations. The results obtained were consistent with the literature with some exceptions such as the H677R (English) and D678N (Tottori) mutation, which were reported to only enhance aggregation and not Aβ peptide accumulation (19).

**Fig 3:**
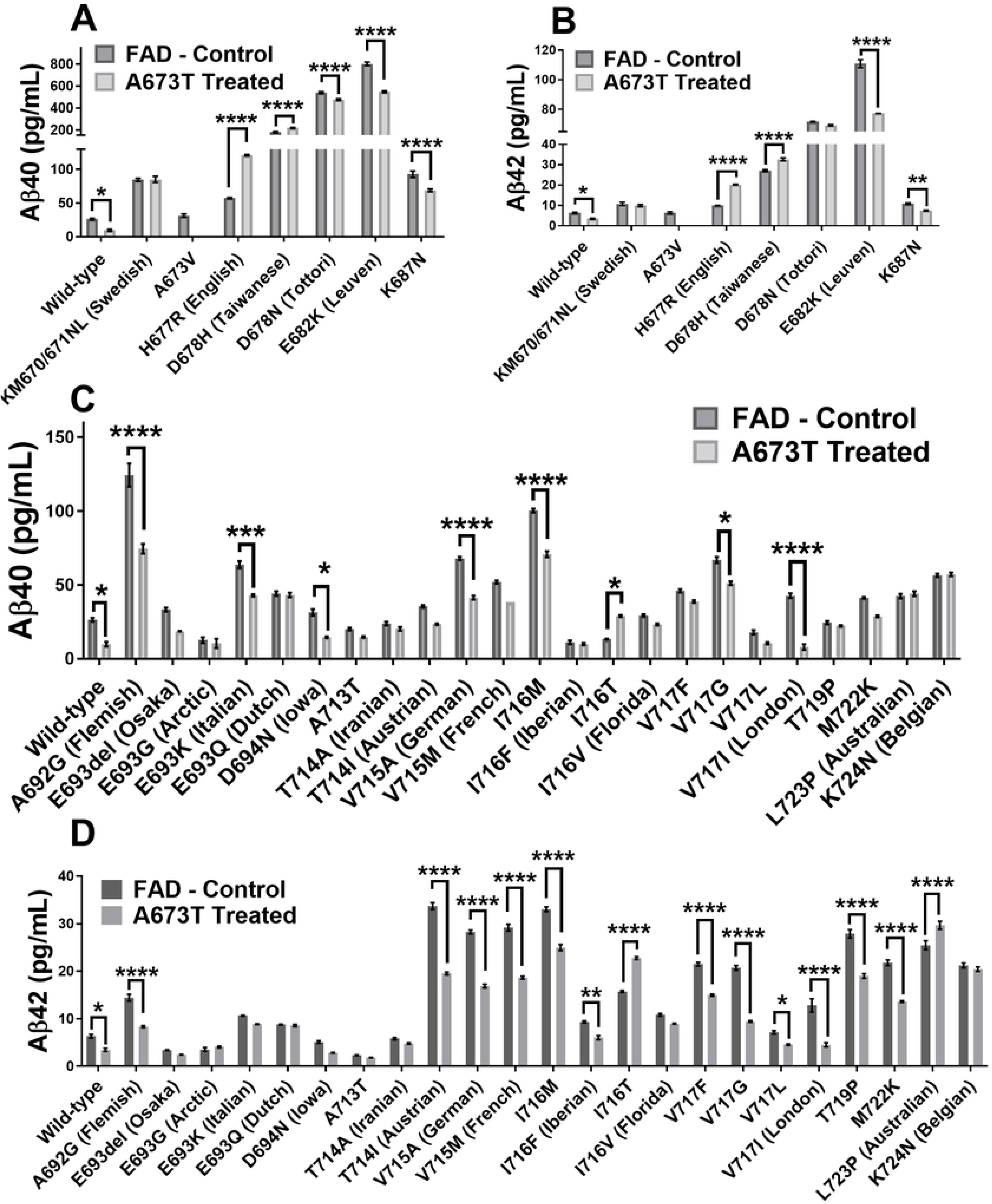
Aβ peptide concentrations in culture medium. Neuroblastomas were transfected with plasmids coding for various FAD mutations with or without an additional Icelandic (A673T) mutation. Exon 16 FAD mutations Aβ40 concentration (**A**) and Aβ42 concentration (**B**). Exon 17 FAD mutations Aβ40 concentration (**C**) and Aβ42 concentration (**D**). Statistic test: Two-way ANOVA Sidak’s multiple comparisons test (n=6). P value style * p<0.0332, ** p<0.0021, *** p<0.0002, **** p<0.0001.

The reduction of Aβ40 and Aβ42 peptide production by the insertion of the additional A673T Islandic mutation is clear for the wild-type APP control gene but also for several FAD mutations. It was further shown that the A673T mutation has a greater effect against FAD mutations in exon 17 than exon 16. The addition of the A673T mutation decreased the Aβ40 concentration for 23 out of 29 FAD plasmids (79%). In addition, Aβ42 concentrations were also decreased in 24 out of 29 FAD plasmids (83%). However, the addition of the A673T mutation increased the Aβ40 concentrations for 6 FAD plasmids (21%) and the Aβ42 concentrations for 5 FAD plasmids (17%) FAD mutations. The Sidak’s multiple comparisons test confirmed a significant Aβ40 reduction for 10 (34%) FAD mutations and a Aβ42 reduction for 14 (48%) FAD mutation. However, for 48% of the FAD mutations, the concentrations of both Aβ40 and Aβ42 peptides were reduced by at least 20% (Fig 4) but not to a wild-type APP gene carrier level.

**Fig 4:**
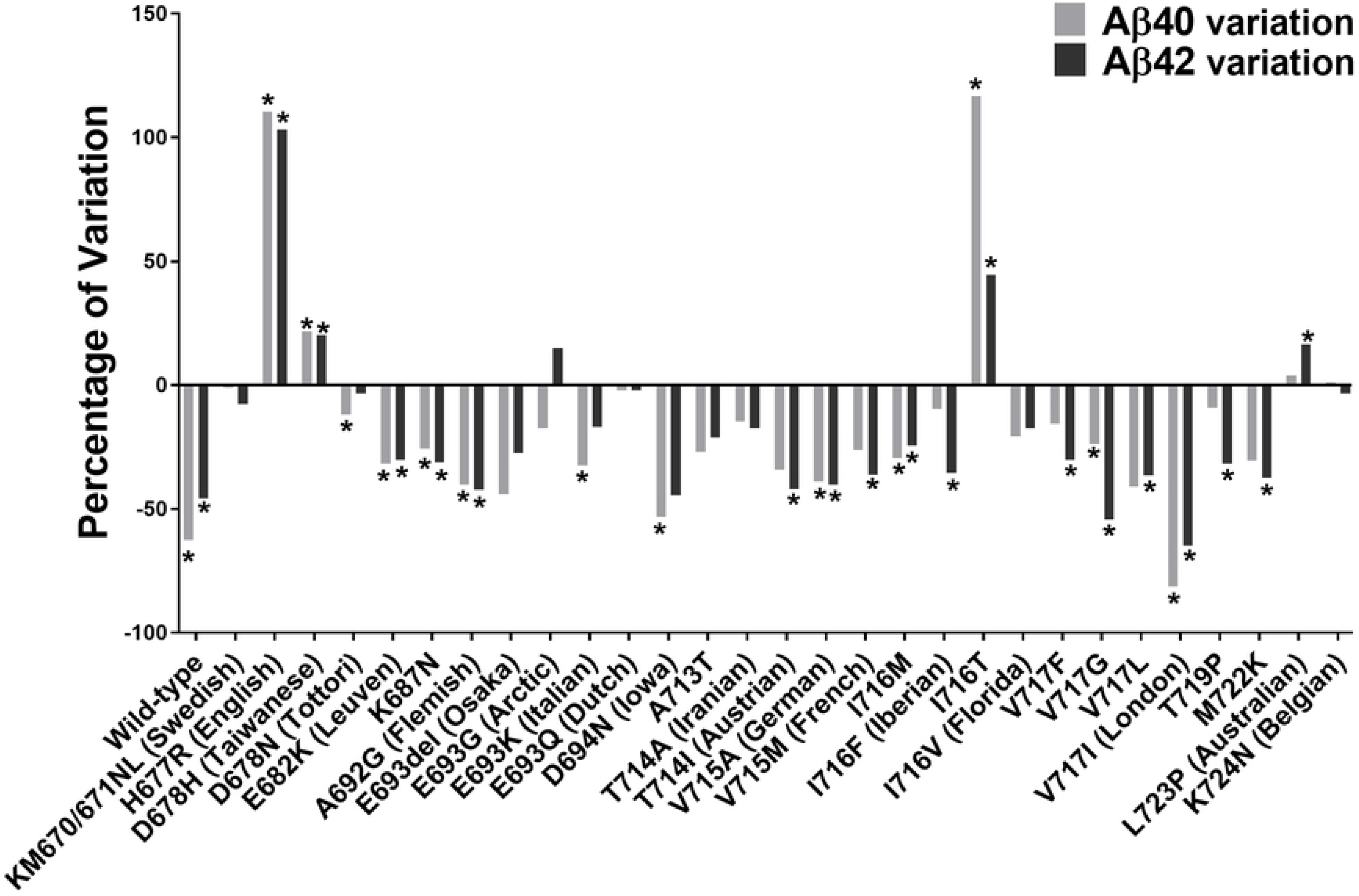
Variations of Aβ40 and Aβ42 peptide concentrations with A673T mutation. Percentage changes of Aβ40 and Aβ42 peptide concentrations induced by the addition of the A673T mutation to plasmids containing or not an FAD mutation. **\*** indicates this variation was statistically significant in the two-way ANOVA Sidak’s multiple comparisons test.

### Aβ42/Aβ40 ratio

The Aβ42/Aβ40 ratios were calculated for all FAD mutations with and without the additional Icelandic mutation (Fig 5). However, since the Icelandic mutation also decreased the Aβ40 concentration, sometimes this ratio was increased by the Icelandic mutation insertion even though both Aβ peptide concentrations were significantly reduced. This was particularly the case for the London (V717I) mutation (20). We thus ranked the different FAD mutations by their capacity to reduce the concentration of Aβ42 in Table 1. Such a ranking will be useful for choosing the right mouse model or patients should one attempt to design an FAD-specific therapy.

**Fig 5:**
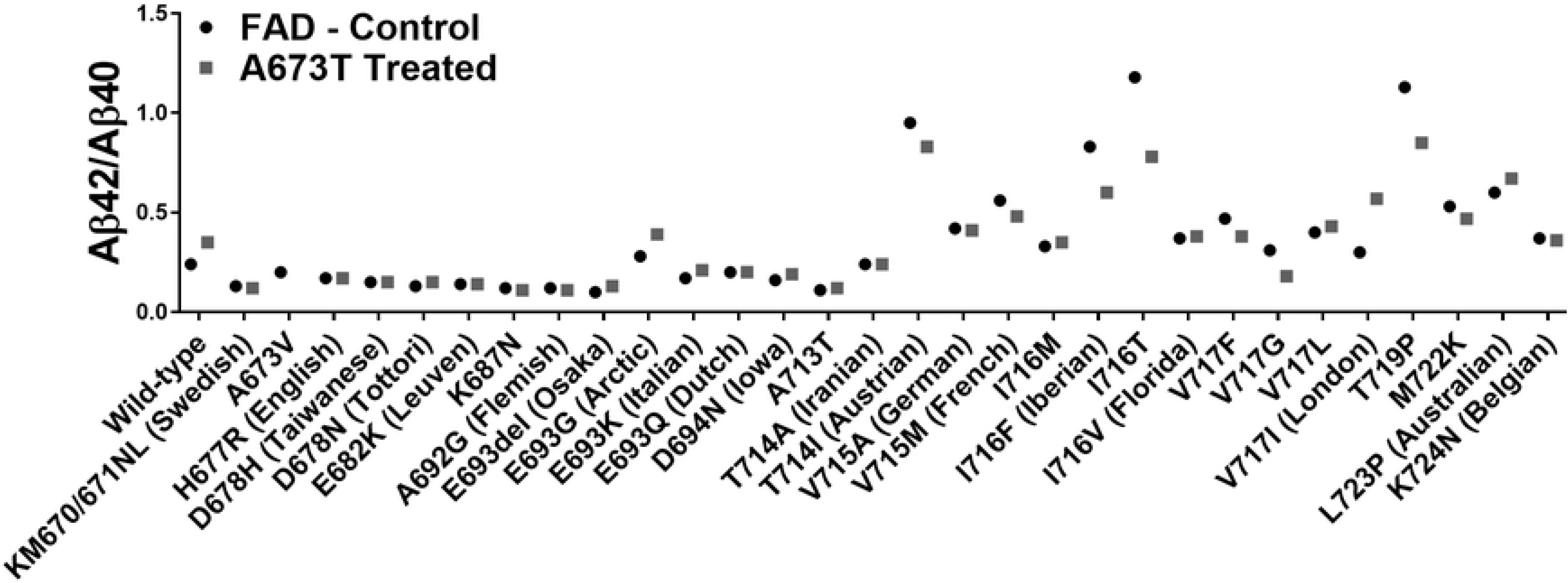
Aβ42/Aβ40 ratios. Culture medium of cells transfected with plasmids coding for various FAD mutations with or without the additional A673T mutation.

**Table 1:** Percentage variations in Aβ40 and Aβ42 peptide concentrations due to the A673T insertion.

| **EOAD mutation** | **Aβ42**<br>**Concentration**<br>**(%)** | **Significancy** | **Aβ40**<br>**Concentration**<br>**(%)** | **Significancy** | **Exon** |
| --- | --- | --- | --- | --- | --- |
| V717I (London) | -65 | **** | -81 | **** | 17 |
| V717G | -54 | **** | -24 | * | 17 |
| Wild-type | -46 | * | -63 | * | na |
| D694N (Iowa) | -44 | ns | -53 | * | 17 |
| A692G (Flemish) | -42 | **** | -40 | **** | 17 |
| T714I (Austrian) | -42 | **** | -34 | ns | 17 |
| V715A (German) | -40 | **** | -39 | **** | 17 |
| M722K | -37 | **** | -30 | ns | 17 |
| V717L | -36 | * | -41 | ns | 17 |
| V715M (French) | -36 | **** | -26 | ns | 17 |
| I716F (Iberian) | -36 | ** | -10 | ns | 17 |
| T719P | -32 | **** | -9 | ns | 17 |
| K687N | -31 | ** | -26 | **** | 16 |
| V717F | -30 | **** | -16 | ns | 17 |
| E682K (Leuven) | -30 | **** | -32 | **** | 16 |
| E693del (Osaka) | -28 | ns | -44 | ns | 17 |
| I716M | -24 | **** | -29 | **** | 17 |
| A713T | -21 | ns | -27 | ns | 17 |
| I716V (Florida) | -17 | ns | -21 | ns | 17 |
| T714A (Iranian) | -17 | ns | -15 | ns | 17 |
| E693K (Italian) | -17 | ns | -32 | *** | 17 |
| KM670/671NL (Swedish) | -8 | ns | 1 | ns | 16 |
| K724N (Belgian) | -3 | ns | 1 | ns | 17 |
| D678N (Tottori) | -3 | ns | -12 | **** | 16 |
| E693Q (Dutch) | -2 | ns | -2 | ns | 17 |
| E693G (Arctic) | 15 | ns | -17 | ns | 17 |
| L723P (Australian) | 16 | **** | 4 | ns | 17 |
| D678H (Taiwanese) | 20 | **** | 22 | **** | 16 |
| I716T | 44 | **** | 117 | * | 17 |
| H677R (English) | 103 | **** | 110 | **** | 16 |

FAD mutations were ranked from the highest decrease in Aβ42 concentration to the highest increase. The significance was determined by a Sidak's ANOVA test. P value style \* p<0.0332, \*\* p<0.0021, \*\*\* p<0.0002, \*\*\*\* p<0.0001.

### The most therapeutically relevant mutation

Among all the FAD mutations of the APP gene, the V717I (London mutation) presented very interesting results following the insertion of the A673T mutation. This mutation was able to diminish the Aβ42 percentage in the extracellular environment by 63% and Aβ40 by 80% (Table 1). The Aβ42 and Aβ40 peptide concentrations were actually reduced to levels that were similar to those observed in the A673T carrier without an FAD mutation (Fig 6).

**Fig 6:**
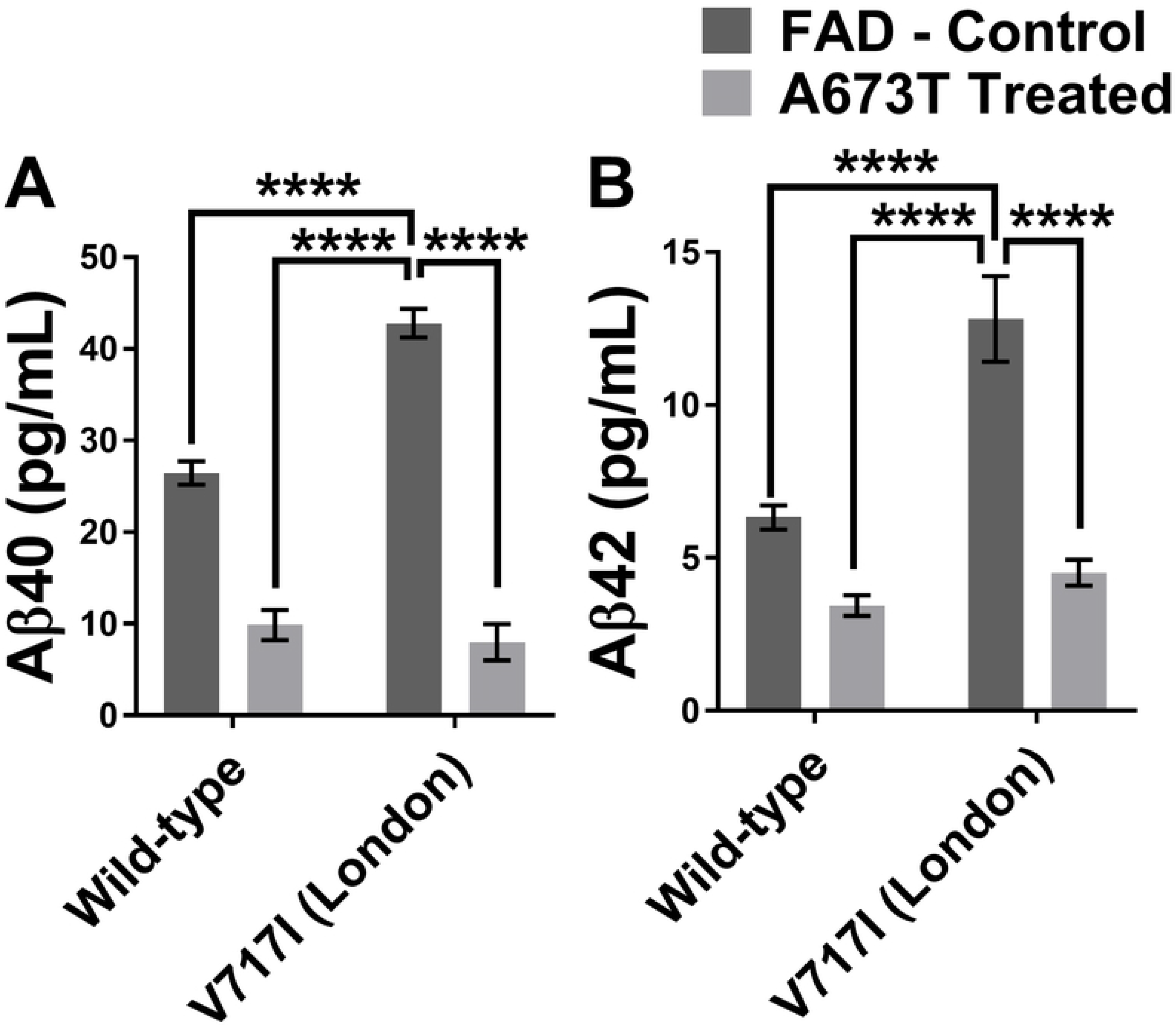
Aβ peptide concentrations with wild type or London APP gene with or without A673T mutation. Aβ40 (in **A**) and Aβ42 (in **B**) concentrations for neuroblastomas transfected with plasmids coding either for a wild type APP gene or an APP gene with the London mutation (V717I) with and without the Icelandic (A673T) mutation. Statistic test: Two-way ANOVA Sidak’s multiple comparisons test (n=6). P value style * p<0.0332, ** p<0.0021, *** p<0.0002, **** p<0.0001.

## Discussion

The A673T mutation has been theorized to provide protective effects against AD onset and development. However, until now, the effects of this mutation on FAD mutations have never been tested in practice. Until now patients with a family history of Alzheimer’s disease have been stuck without an effective treatment to prevent the loss of their mental faculties starting as early as their late 40’s (21). Here, we showed that the addition of the A673T mutation to an APP cDNA containing an FAD mutation can decrease the secretion of Aβ42 and Aβ40 peptides for most FAD mutations.

One notable observation in this study was that the A673T mutation generally had stronger protective effects against FAD mutations in exon 17 compared to exon 16. This may be because some mutations in exon 16 are located close to the BACE1 cutting site and interfere with the protective effect of the A673T mutation, which reduces cutting of the APP protein. The additional FAD mutation may be favoring a structural conformation of the APP protein which favors cleavage by BACE1, thereby increasing Aβ40 and Aβ42 peptide accumulation.

Another interesting observation of this study was the drastic difference in the presence of Aβ peptides, especially when different mutations occurred coding for the same amino acid. It was noted that A673T has a strong protective effect on the I716M mutation and correspondingly reduced the formation of both Aβ40 and Aβ42 peptides. In A716T however, the presence of A673T actually increased the formation of both peptides.

Indeed, for some FAD mutations, the addition of the A673T mutation resulted in an increase rather than a decrease in the concentration of Aβ peptides (Fig 4). Five FAD mutations seemed to increase in severity of Aβ peptide accumulation. However, an increase of Aβ40 may not necessarily accentuate the severity of the Alzheimer’s disease since it is the Aβ42/ Aβ40 ratio which determines aggregation and subsequent disease onset/development. However, for the purposes of a treatment, a clear reduction of both Aβ peptides is likely beneficial.

The reduction of Aβ peptides in the extracellular environment following the introduction of the A673T mutation was especially encouraging in the case of the V717I London mutation. Not only did the percentages of Aβ42 and Aβ40 demonstrate the greatest reductions with this mutation (Table 1) but the London mutation is also one of the most common FAD mutations in the world (ALZ.org). Our results demonstrate that the addition of the A673T mutation in a London patient has the potential to reduce their Aβ peptide levels to that of healthy AD-free individuals (Fig 6) and serve as a promising avenue for a gene therapy. It would probably be difficult or even impossible to obtain ethical approval for a Phase I clinical trial for sporadic Alzheimer patients in a preclinical state, i.e., before symptom development. However, the data observed and discussed in this article stands to help validate the launch of Phase I clinical trials for this kind of gene therapy for London patients around the world (around 30 families). It would make the development of a Phase I clinical trial much simpler since we would be able to know the genotype of the patients several years before the symptom apparition. Down the line, these findings may also serve to validate the inception of the same gene therapy for sporadic AD patients. Our results also support the development of other gene therapies that are predicated on the addition of a protective mutation as opposed to the return of the gene to the wildtype.

In certain cases, the percentage decrease of Aβ40 and Aβ42 is insufficient to determine whether an FAD mutation is an optimal candidate for A673T treatment. For example, the Leuven E682K mutation demonstrated a considerable reduction of Aβ40 and Aβ42 concentrations (Table 1). However, this diminution was incapable of bringing the peptide concentrations to acceptable levels. Rather, the peptides were still present in a concentration that was severalfold that of the wildtype APP gene (Fig 3A and 3B). Ultimately, a gene therapy for this FAD mutation using A673T may slow down the progression of the disease as the concentration of the Aβ peptides is diminished; however, the data does not suggest that this treatment alone has the potential to prevent the onset of and development of AD based on Aβ peptide secretion.

Our report has solely studied the secretion of the Aβ peptides. It must be remembered that the aggregation of these peptides is also a very important parameter as it plays an essential role in the protective effect of A673T (22). Most FAD mutations are pathogenic due to the changes in the aggregation of the Aβ peptides as a result of the amino acid modifications. Adding A673T, which is also known to reduce aggregation, may create some competition with the pathogenic mutation and reduce aggregation but the results are hard to predict without direct experimentation (22). It is possible that some FAD mutations showing only a moderate reduction of Aβ peptide production following the insertion of the A673T mutation may experience a significant reduction in overall aggregation of said peptides.

Our next step will be to test the potential protective effect of A673T *in vitro* and *in vivo* by inserting the A673T mutation directly in cells derived from FAD patients or in mouse models using base editing (23) or PRIME editing (24). This will serve as a proof of concept in the development of a gene therapy based on the A673T Icelandic mutation insertion. We raise the hypothesis that this approach will allow clinicians to eventually treat a large number of persons affected by a sporadic AD. It has already been proven that the A673T mutation protects the natural carrier of this mutation, so we suggest that the artificial insertion of the mutation could help most sporadic AD cases as well. We are proposing that the dominant A673T protective mutation may compensate for most genetic risk factors.

This project has demonstrated that the insertion of the A673T mutation is beneficial for patients with most forms of FAD mutations in the APP gene. Our future experiments will attempt to verify whether the A673T mutation in APP is also protective in trans for other genes that have been related to AD such as the PSEN1 or PSEN2.

## Methods

### Construction of plasmid libraries containing an FAD mutation

The backbone plasmid pcDNA6/V5-His was purchased from Invitrogen Inc. (Carlsbad, CA). The APP695 cDNA (courtesy of Dr. G. Levesque, CHUQ, Quebec) was inserted by ligation between Kpn1 and Xba1 cut sites. The ensuing plasmid was then mutated using the New England Biolabs (NEB, Ipswich, MA) mutagenesis Q5 kit in 29 different reactions. The 29 new plasmids each represented a form of FAD and served as a “normal” library version of each FAD. The mutations were located in exons 16 and 17 to better demonstrate the protective effects of A673T. Another “mutated” library was created by adding an additional A673T mutation to each FAD plasmid. Prior to the start of the experiments, the plasmids underwent Sanger sequencing to ensure that the only mutations present were those under study.

### Transfection in SH-SY5Y of plasmid libraries

The transfection reagent (Lipofectamine 2000TM) and Opti-MEM-1™ culture media were purchased from Life Technologies Inc. (Carlsbad, CA). The day before the transfection, 100,000 SH-SY5Y cells were seeded per well in 24 well plates in DMEM/F12 supplemented with 10% Fetal Bovine Serum (FBS) and antibiotics (penicillin/streptomycin 100 μg/mL). The following morning, the culture medium was changed for 500 μl of DMEM/F12 medium supplemented with 10% FBS without antibiotics. The plate was maintained at 37°C for the time required to prepare the transfection solution. For the transfection, solutions A and B were first prepared. Solution A contained 48 μl of Opti-MEM-1™ and 2 μl of Lipofectamine™ 2000 for a final volume of 50 μl. Solution B was prepared as follows: a volume of DNA solution containing 800 ng of DNA was mixed with a volume of Opti-MEM-1™ to obtain a final volume of 50 μl. Solutions A and B were then mixed together by up and down movements and incubated at room temperature for 20 minutes. 100 μl of the ensuing solution were then added to each well. The plate was left in the CO_2_ incubator for a period of 4 to 6 hours. The medium was replaced by 500 μl of DMEM F12 supplemented with 10% FBS and antibiotics. The plate was kept for 72 hours in the CO_2_ incubator before extraction of genomic DNA. The culture medium was harvested and protease inhibitors (1 mM PMSF + 1X complete tabs from Roche) were added. The media were then stored at −80°C.

### Culture medium analysis

The concentrations of Aβ40 and Aβ42 peptides were measured with Meso Scale Discovery Inc. (MSD, Rockville, MA) Neurodegenerative Disease Assay 6E10 kit. Standards and samples were prepared according to the manufacturer’s protocols and tested in triplicate for each experiment.

### Statistical analysis

All statistic tests and graphs were performed as recommended by GraphPad Prism 7.0. Two-way ANOVA Sidak’s multiple comparisons test was used to test significance with three biological replicas (two technical replicas each) for Fig 2, 3 and 6. P value style * p<0.0332, ** p<0.0021, *** p<0.0002, **** p<0.0001.

## Acknowledgment

We would like to thank Dr. G. Levesque (CHUQ, Quebec) for giving us the APP695 cDNA plasmid as well as Dr R. Lapointe (CHUM, Montréal) for allowing us to use the MSD equipment in his laboratory.

## Supporting information

**S1_data**

